# Power-law memory governs bacterial adaptation and learning in fluctuating environments

**DOI:** 10.1101/2025.04.26.650739

**Authors:** Josiah C. Kratz, Huijing Wang, Fangwei Si, Shiladitya Banerjee

**Affiliations:** Computational Biology Department, Carnegie Mellon University, Pittsburgh, PA 15213, USA; Department of Physics, Carnegie Mellon University, Pittsburgh, PA 15213, USA; School of Physics, Georgia Institute of Technology, Atlanta, GA 30332, USA

## Abstract

How do single-celled organisms adapt and learn to survive in dynamic environments without a nervous system? Here, we provide experimental evidence and a theoretical model demonstrating learning-like behavior by single bacterial cells in fluctuating environments. Using a custom microfluidic platform, we tracked individual *E. coli* cells in dynamic nutrient environments and found that bacteria adapt on multiple timescales, tuning their growth control behavior based on prior environmental experience. Motivated by our observation that cellular adaptation dynamics are approximately scale-free, we built a theoretical framework for bacterial growth control with dynamic power-law memory to explain how bacteria integrate environmental information over a range of timescales to enable growth rate adaptation. We show how this behavior arises naturally from heterogeneous ribosomal relaxation dynamics within a bacterial cell. Using this model, we identify an inherent trade-off between growth rate maximization and adaptation speed, which we validate experimentally in pulsatile nutrient environments. Finally, we connect our mechanistic reaction-network model to descriptions of artificial recurrent neural networks, identifying a minimal network architecture capable of exhibiting adaptation and learning at the single-cell level.

---

Single-celled microorganisms exhibit remarkable phenotypic plasticity, dynamically reshaping their intracellular composition to adapt to diverse conditions (1–4). To do so, cells utilize environmental cues to modulate gene expression, leading to condition-dependent changes in physiology and behavior (5–8). Remarkably, such complex responses emerge in the absence of any nervous system, yet allow microbes to thrive under a wide range of stressors (9–14). While much progress has been made in linking environmental inputs to gene expression (15–19), it remains unclear whether single cells can learn from past experiences —integrating temporal information to guide future behavior. Resolving this question represents a key challenge in identifying the physical principles that govern adaptation, memory, and decision-making at the cellular level.

Bacterial cells are known to exhibit adaptive behaviors in fluctuating environments, where prior environmental experiences can shape future responses (5, 7, 8, 20, 21). These include memory-like traits such as faster adaptation to repeated nutrient or antibiotic shifts (7, 8). Specifically, it has been shown that bacteria exhibit faster cellular response times in fluctuating nutrient environments, but do so at the expense of growth (Fig. 1a) (7). Such trade-offs represent a fundamental balance between responsiveness and fitness. While theoretical models have explored how environmental stress profiles influence phenotypic switching and resource allocation strategies (4, 15, 16, 18, 22–26), they typically assume single-timescale dynamics and lack mechanisms for integrating environmental history into cellular decision-making. As a result, the physical principles by which single cells adaptively tune their growth in dynamic environments remain poorly understood.

**Fig. 1.**
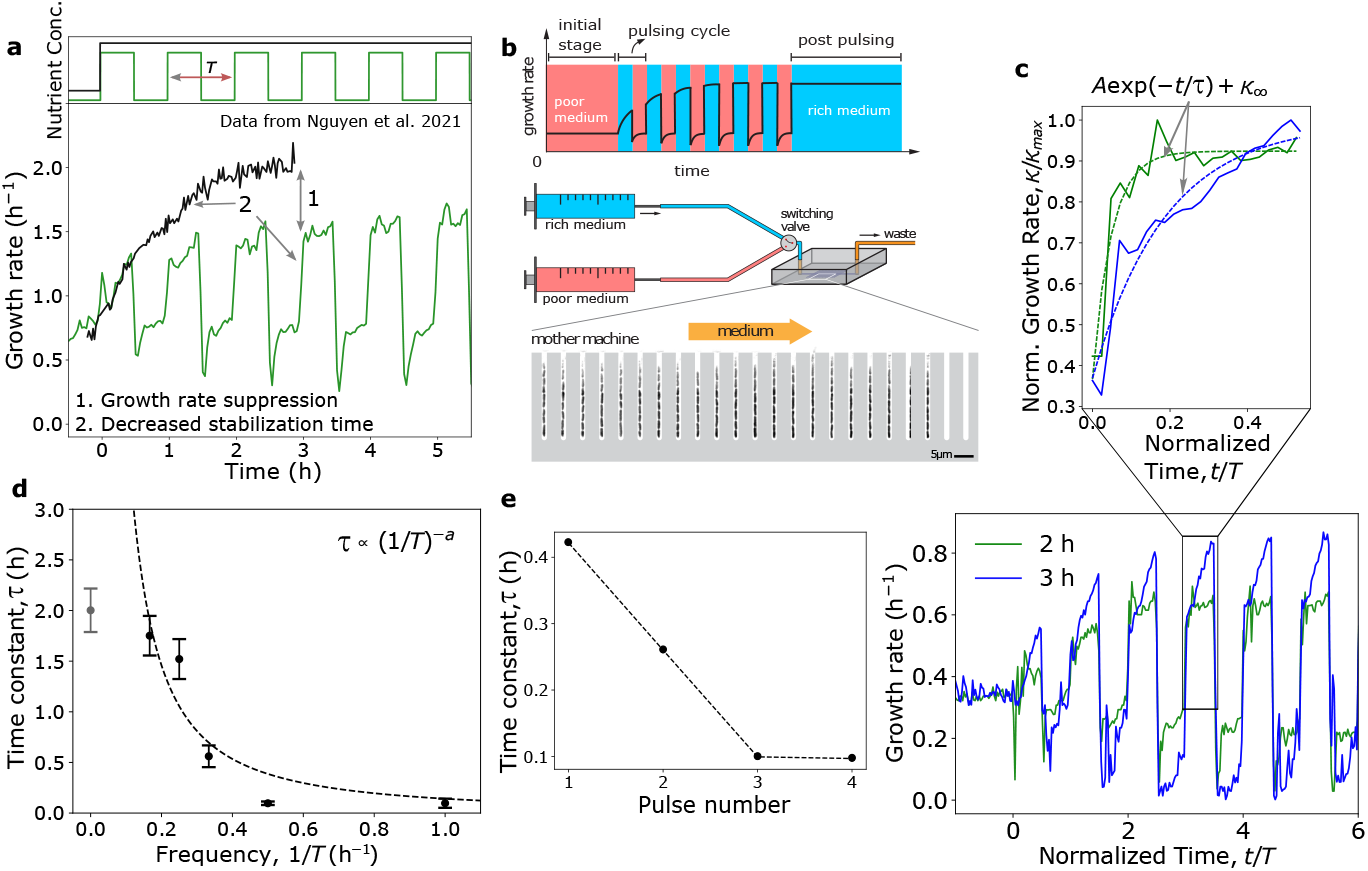
History-dependent adaptation in bacterial growth control. **a** In contrast to bacteria experiencing a single nutrient upshift (black), bacteria subjected to pulsatile nutrient-rich pulses (green) increase growth rate following upshift significantly faster after experiencing several pulses, but growth rate stabilizes at a value well below that of cells experiencing constant nutrient-rich conditions. Experimental data are from ref. (7) and are of *E. coli* K-12 NCM3722 *ΔmotA* cells switching between low (0.1%) and high (2%) concentrations of LB. **b** To elucidate the dependence of growth rate behavior on the time-profile of nutrient fluctuations, we measured single-cell growth rate of *E. coli* K-12 MG1655 cells subjected to nutrient shifts between MOPS sodium acetate and MOPS glucose with amino acids using a custom microfluidic device (see Methods) for both single shifts and for pulses over a range of time periods, *T*. **c, bottom** Two example trajectories are shown for pulsing with a time period of *T* = 2 h (purple) and *T* = 3 h (green). For each condition, single-cell growth rates were calculated from volume trajectories and averaged across the population through time (see Methods), resulting in ∼30,000 cells being tracked over the course of each experiment (see Supplementary Table 3 for list of experimental conditions and sample size). **c, top** For each pulsing frequency, the effective adaptation time constant, *τ*, was obtained by fitting the first upshift after 6 h to the exponential *A* exp(−*t/τ*) + *κ*_*∞*_ (best fit shown by the dashed line for each example). **d** Adaptation time constant *τ* decreases with increasing pulse frequency */T*. The time constants were obtained from the fitting procedure described in **c**, where error bars indicate uncertainty in the exponential fit. Note that a frequency of 0 corresponds to a single nutrient upshift (purple circle). This point was omitted from fitting the power law relating *T* and *τ* (dashed line). **e** For cells initially grown in nutrient-poor media as in **c**, adaptation time constant decreases with subsequent nutrient-rich pulses before stabilizing. Time constants shown are for *T* = 2 h, and are obtained from the fitting procedure described in **c**.

In this study, we use *Escherichia coli* as a model organism to investigate learning-like behavior at the single-cell level. Using a high-throughput microfluidic platform, we track individual *E. coli* cells exposed to pulsatile nutrient signals and show that these cells are capable of adapting over multiple timescales, tuning their growth control strategies based on environmental patterns. Our data reveal that the timescale of adaptation decreases as the frequency of nutrient fluctuations increases, but this faster response comes at the expense of reduced growth rates. We show that this behavior stems from a power-law memory in proteome allocation, driven by heterogeneous ribosomal relaxation dynamics, providing a mechanistic model of bacterial memory and continual learning. This framework captures key features of bacterial learning and allows us to map bacterial adaptation onto the dynamics of artificial recurrent neural networks, identifying the minimal architecture required for history-dependent behavior at the single-cell level. Together, our results establish a conceptual and quantitative foundation for understanding learning in living cells, revealing how even the simplest organisms can integrate past experience to navigate dynamic environments.

## Bacterial cells exhibit multiple timescales of adaptation in fluctuating nutrient environments

To quantitatively characterize how the time-profile of nutrient fluctuations affects growth control and adaptation, we subjected *E. coli* cells to precisely controlled nutrient shifts between a nutrient-poor medium (MOPS Sodium acetate) and a nutrient-rich medium (MOPS glucose with 11 amino acids) in a mother machine (Fig. 1b, see Methods for experimental details). We measured single-cell growth rate dynamics in response to nutrient oscillations with different time periods *T* (Fig. 1c, bottom). For each frequency *T* ^*−*1^, we obtained the timescale *τ* of growth adaptation by fitting the average growth rate trajectory to an exponential of the form *A* exp(−*t/τ*) + *κ*_*∞*_ (Fig. 1c, top). For cells initially grown in nutrient-poor media, this timescale decreased with subsequent nutrient-rich pulses, indicative of episodic learning (Fig. 1e). In addition, our data indicate that *τ* scales approximately as a power law with *T*, showing bacteria exhibit multiple timescales of adaptation and suggesting scale-free behavior (Fig. 1d). These data indicate that single cells integrate environmental information over both short and long durations to facilitate rapid adaptation.

To understand the mechanisms underpinning memory and adaptation in bacterial growth, we tested an existing model of bacterial growth control based on proteome allocation theory, which has previously been validated in dynamic environments (18, 24, 27). In this framework, the environment is defined by the nutrient availability *c*(*t*), which we define as the product of nutrient concentration and quality (yield). The cell imports nutrients and converts them into amino acids with mass fraction *a* by metabolic proteins. Amino acids are consumed by translating ribosomes, to synthesize each of the three proteome sectors defined by their mass fraction: ribosomal (*ϕ*_*R*_), metabolic (*ϕ*_*P*_), and “housekeeping” (*ϕ*_*Q*_) (Fig. 2a). The ribosomal mass fraction sets the exponential growth rate, given by

**Fig. 2.**
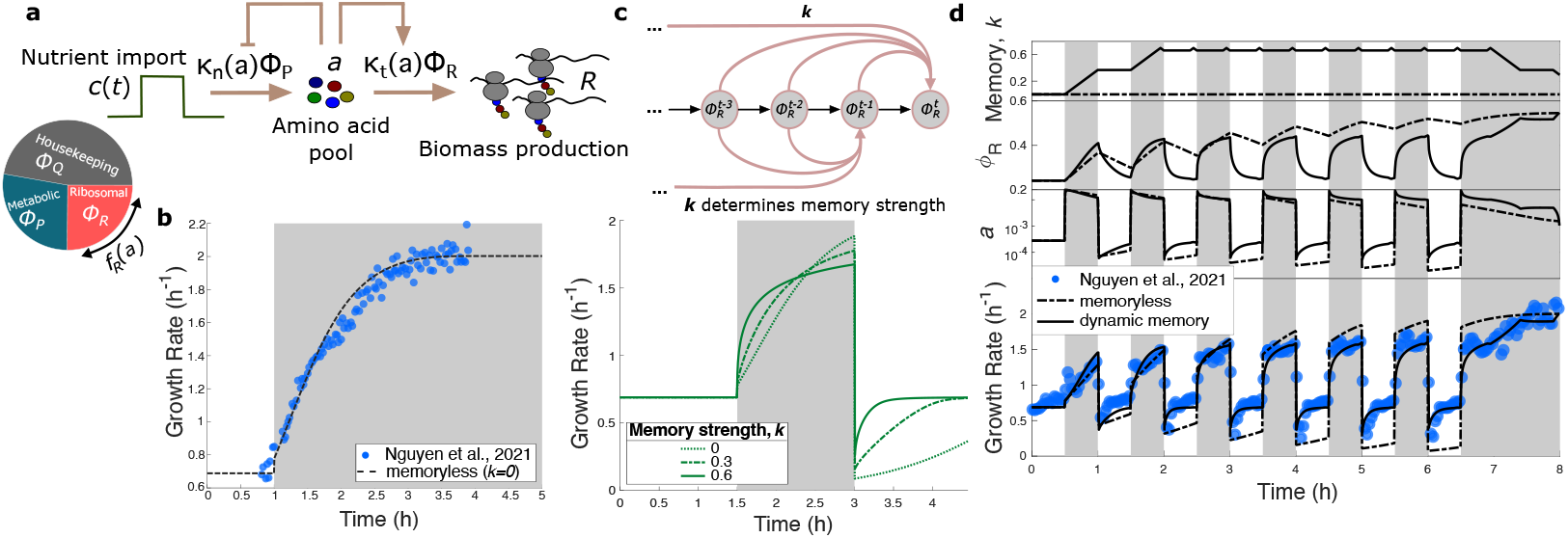
Dynamic memory model of bacterial growth control. **a** Schematic of coarse-grained model of bacterial growth control. Nutrients (c) are imported by metabolic proteins (P) and converted to amino acids (a), which are then consumed by ribosomes (R) to biomass. **b** Simulation dynamics of growth rate control in response to nutrient upshift for the memoryless model. 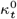 was obtained by fitting directly to growth rate data. **c** (top) Schematic depicting the time nonlocality caused by introduction of a power law memory kernel. The ribosomal dynamics at time *t* are now a function of all previous values. The strength of the influence of past values on the current time is controlled by the memory strength *k*. (bottom) Growth rate dynamics in response to a pulse of nutrients for different constant memory strength values. Parameter values are identical to those in (**b**) except *k*. **d** Corresponding memory strength (top), ribosomal mass fraction (2nd from top), amino acid mass fraction (2nd from bottom), and growth rate (bottom) dynamics for both the memoryless (dashed) and the dynamic memory model (solid). To match with experiment, cells were initialized in poor nutrient conditions followed by nutrient-rich pulsing. While the memoryless model is unable to capture observed behavior, the dynamic memory model agrees well with experimental data. For **b, c**, and **d**, nutrient-rich periods are denoted by gray shading. All experimental data are from ref. (7). See Table 1 in Supplementary Information for a list of parameters.

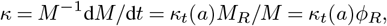

where *M* is the total protein mass of the cell, *M*_*R*_ is the ribosomal mass and *κ*_*t*_(*a*) is the translational efficiency (see Supplementary Note 1 for details). The dynamics for the ribosomal sector can be written as:

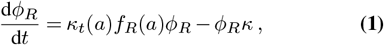

where *f*_*R*_(*a*) denotes the fraction of total synthesis capacity devoted to ribosomes, and is chosen to maximize translational flux at steady state, thus maximizing growth rate (16, 27–29). A significant portion of the *E. coli* proteome is invariant to environmental perturbations (2, 3, 6), thus we take the housekeeping sector to be constant, such that *ϕ*_*Q*_ = *f*_*Q*_ = const. and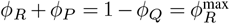. This allows the amino acid dynamics, which are coupled to Eq. (1), to be written as:

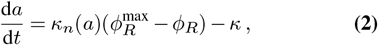

where *κ*_*n*_(*a*) is the nutritional efficiency (see Supplementary Note 1 for details). In this way, *a* acts as a read-out of flux imbalance, and so by altering proteome allocation in response to *a*, the cell can dynamically respond to nutrient changes. In this model, there exists a unique proteome allocation (value of *ϕ*_*R*_) which maximizes growth in a given nutrient environment (value of *c*). Thus, in response to an environmental change, bacteria can adapt by reshaping their proteome to increase fitness. This requires global reallocation of protein synthesis (change in *f*_*R*_), as well as growth-mediated dilution of the current suboptimal proteome – a process which in this model occurs over a single timescale set by the translational efficiency *κ*_*t*_. This framework can accurately capture the ribosomal growth law (30) when evaluated at steady-state (constant nutrient environment), as well as single nutrient shifts (Fig. 2b). However, this minimal model fails to capture experimentally observed growth control in pulsatile environments (Fig. 2d). Most notably, growth rate fails to recover following each nutrient downshift, and after several pulses growth rate exceeds experimentally measured values. Furthermore, the predicted adaptation timescale shows no correlation with the time period of the nutrient pulse (Fig. 3d), contrary to experimental data.

**Fig. 3.**
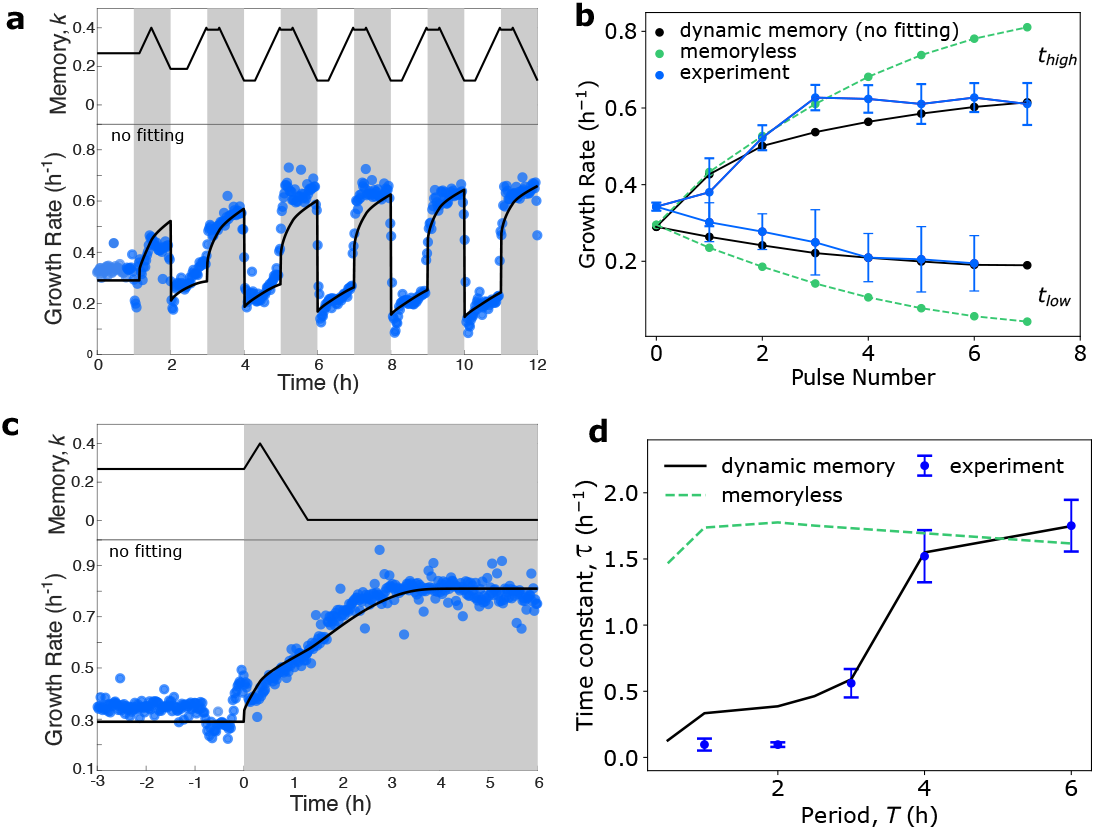
Dynamic memory model captures wide range of observed growth control behavior. **a** Predicted memory strength dynamics (top) and corresponding growth rate dynamics (bottom) for the proposed dynamic power-law memory model for cells initialized in nutrient-poor conditions followed by nutrient-rich pulsing, compared to our experimental growth rate measurements. **b** Average growth rate during the nutrient-rich half-periods (*t*_high_) and the nutrient-poor half-periods (*t*_low_) for cells initialized in poor nutrient conditions (period 0) followed by nutrient-rich pulsing for the dynamic memory model (quantification of growth rate dynamics shown in **a**) compared to the memoryless model and experimental data. Error bars represent standard deviation of the average growth rate. **c** Predicted memory strength dynamics (top) and corresponding growth rate dynamics (bottom) in response to nutrient upshift for the dynamic memory model, compared to our experimental growth rate measurements. Parameters are identical to that of **a. d** Effective adaptation time constant, *τ*, as a function of time period of nutrient pulsing, *T*, for both the dynamic memory and memoryless models. Model parameters were obtained by fitting to the data in **d**, and then were used to make growth rate predictions for simulations in **a**-**c**. Procedure for obtaining *τ* is identical to that described in Fig. 1d, where error bars indicate uncertainty in the exponential fit. In contrast to Fig. 2d, a bias for rich nutrients (*α*, see Methods for more details) is included, causing *k >* 0 even when in the constant poor nutrient condition. See Table 1 in Supplementary Information for a list of parameters. For both the pulsatile (**a**) and single shift (**c**) conditions, single-cell growth rates were calculated from volume trajectories and averaged across the population through time (see Methods), resulting in ∼30,000 cells being tracked over the course of each experiment (see Supplementary Table 3 for list of experimental conditions and sample size).

## Dynamic power-law memory governs bacterial response to nutrient pulsing

Our data show that adaptation timescale (*τ*) follows an approximate power law relation with nutrient pulse frequency (1*/T*), indicating that bacterial growth control is governed by a scale-invariant process. Scale-free power law dynamics are reflective of fractional-order differential equations (FDEs, cf. (31, 32)), which characterizes systems that integrate input-output relationships over multiple timescales (33). Motivated by this, we first implement a phenomenological model that incorporates a power-law memory kernel into the dynamics of proteome allocation (Eq. (1)), enabling history-dependent responses across diverse timescales (Fig. 2c). With this addition, the dynamics of *ϕ*_*R*_ now become:

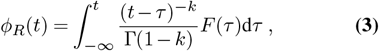

where Γ(·) denotes the Gamma function, *F* (*t*) = *ϕ*_*R*_(*t*)[*κ*_*t*_(*a*(*t*))*f*_*R*_(*a*(*t*)) − *κ*(*t*)] is the total net production-dilution flux given by Eq. (1), and *k ∈* [0, 1) is the power law exponent. Eq. (3) can be written as d^*k*^*ϕ*_*R*_*/*d*t*^*k*^ = *F* (*t*), implying that *k* controls the fractional order, and in turn the degree of nonlocality (memory strength). Thus, a value of *k* = 0 corresponds to the memoryless case (first-order derivative), whereas larger values of *k* result in an increased influence of past states on the current dynamics. The effect of adding a memory kernel to the model can be made clear by analyzing its effect on resulting growth rate trajectories. In response to a single nutrient pulse, including a nonzero memory strength (*k >* 0) results in an initial period of rapid growth rate increase following upshift, but comes at the cost of reduced growth rate at longer timescales compared to the memoryless model (*k* = 0) (Fig. 2c). Conversely, increasing memory strength aids in growth rate recovery following downshift.

While the power-law model with a constant memory strength captures growth rate repression and rapid adaptation behavior of cells stabilized in a pulsatile nutrient environment, it does not capture the result that adaptation rate increases with successive periods of nutrient pulse and the episodic increase in growth seen by cells which are first grown in poor media before being exposed to pulsatile conditions (Supplementary Figs. 1, 2). This suggested that in fluctuating conditions, instead of optimizing for growth rate, bacteria maintain a memory of past environments which allows for increased response time to changing environments. This motivated us to consider a dynamic memory model of growth control, in which *k* is a function of time through its dependence on the environment, *k*(*t*) = *k*(*c*(*t*)), as cells which can tune their memory strength to match the environmental fluctuations are able to thrive in diverse conditions.

To model this cellular decision-making process, we introduced a Bayesian inference scheme (see Methods) in which a cell dynamically updates its estimate of the environment based on previous nutrient signals. This is equivalent to tracking the moving average of nutrient signal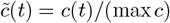, over the interval [*t* − *T*_*N*_, *t*]:

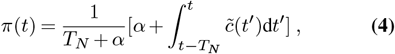

where *α* encodes a bias for rich nutrient environments (see Methods). In Supplementary Note 2 we show how this type of computation could approximately be carried out by the cell. *π*(*t*) serves as the cell’s internal representation of the environment which takes into account historical context, enabling it to accurately discriminate between different environmental time-profiles. In maximally uncertain environments (when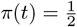), the timescale of growth rate adaptation, and thus proteome reallocation, should be minimized to allow for rapid adaptation to unpredictable environmental changes. Thus we relate the memory strength *k*(*t*) to *π*(*t*) through a simple linear relation: *k*(*t*) *∝* 1 −|2*π*(*t*) − 1|, such that *k* increases in fluctuating environments.

Using this framework, we can simulate the full dynamic memory model to make predictions for bacterial growth control in fluctuating environments with arbitrary pulse frequency (see Methods for numerical simulation details). As seen in Fig. 2d and Fig. 3a, our proposed model is able to successfully capture experimentally-observed *E. coli* growth rate dynamics for cells experiencing a transition from constant to pulsatile nutrient environments with different pulsing frequencies and nutrient types, highlighting the generality of our model and the robustness of the observed growth control phenomena.

Furthermore, for cells initialized in poor nutrients which then experience the onset of rich nutrient pulses, the dynamic memory model predicts an increase in average growth rate in rich nutrients for several pulses before stabilizing at a value well below the steady-state growth rate, matching trends seen in our experimental data (Fig. 3b). In contrast, growth rate converges to the steady-state value in the memoryless model. Additionally, during nutrient-poor half-periods, growth rate in the memoryless model falls to zero, whereas average growth rate in the dynamic memory model stabilizes well above zero, indicative of the quick growth rate recovery caused by the maintenance of a nonzero memory (Fig. 2d and Fig. 3a).

Critically, our model can simultaneously capture single-shift growth rate dynamics with no change in parameter values due to the rapid decay of memory in constant environments (Fig. 3c), underscoring the importance of the dynamic memory formulation. Furthermore, our model captures the observed increase in adaptation timescale with the time period of nutrient oscillation (Fig. 3d), suggesting that the learning traits seen in experiment are a consequence of bacterial regulatory strategies which tune ribosomal adaptation dynamics to match the time-profile of nutrient fluctuations.

## Mechanistic origin of power-law memory

To explain the mechanistic origin of power-law memory present in our model (Eq. (3)), we propose a model of growth control based on heterogeneous ribosomal dynamics in a bacterial cell. Contributions to cell growth arise from many ribosomes that differ in composition (34), and are subject to distinct feedback mechanisms that act over a wide range of timescales in response to environmental stress (35–37). For example, on short timescales (minutes), translation initiation is inhibited by ppGpp, a small molecule whose concentration increases under nutrient limitation (37, 38). On longer timescales (hours), nutrient depletion can induce the expression of hibernation factors (35, 39–41), which bind and inactivate subsets of ribosomes, and ribosomes are also subject to active degradation depending on nutrient availability (42, 43).

Together, these regulatory processes suggest that ribosomes can be grouped into functional sub-populations, or subsectors, each experiencing distinct feedback control and turnover kinetics. These subsectors may reflect differences in the types of mRNAs being translated, initiation efficiencies, or other regulatory features that influence mRNA-specific translation rates and ribosome degradation. Motivated by this, we partition the total ribosome mass fraction, *ϕ*_*R*_, into *m* sub-sectors, each with a characteristic relaxation timescale *λ*_*m*_ (Fig. 4a). This yields the dynamics:

**Fig. 4.**
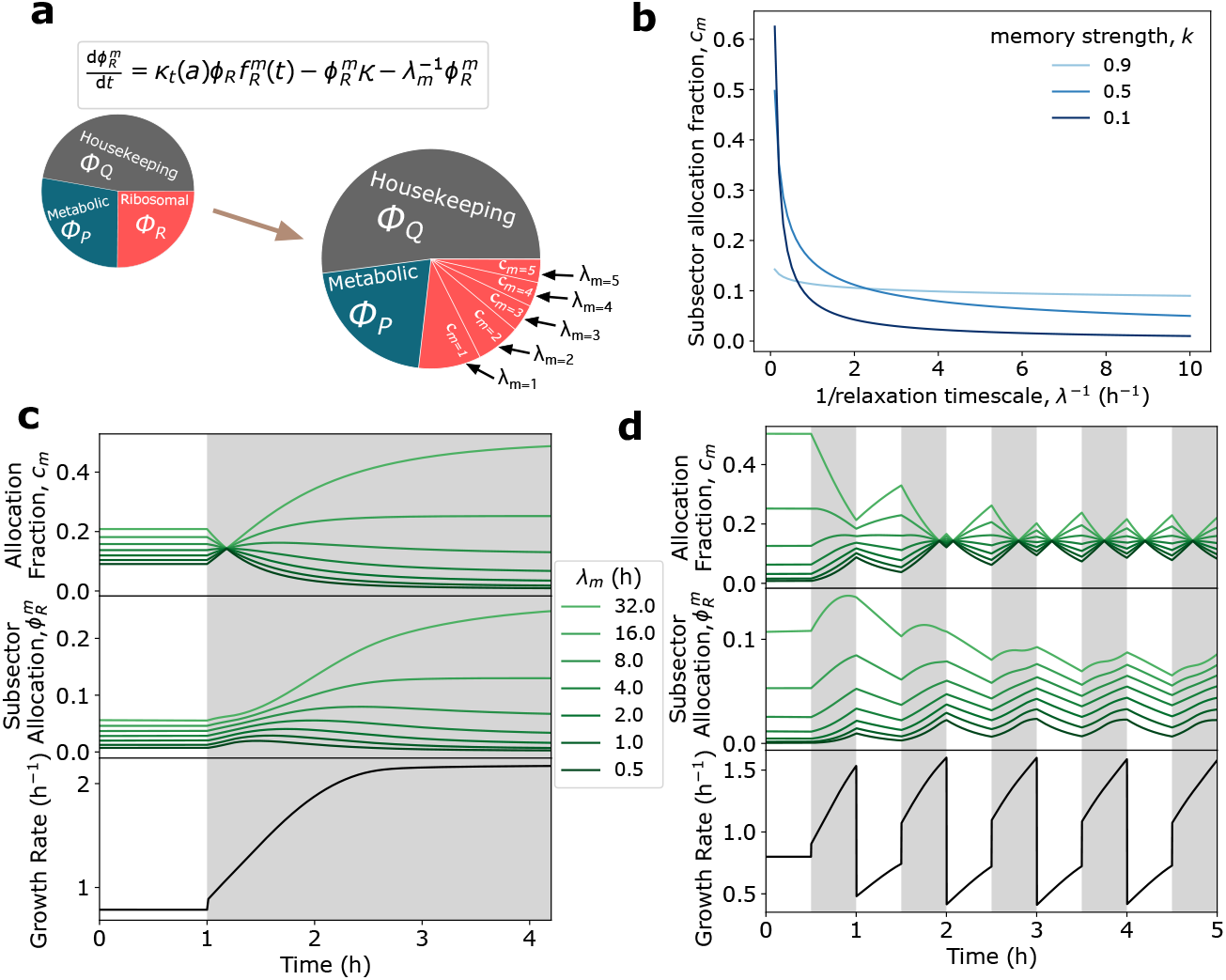
Emergence of a fractional model of growth control from heterogeneous ribosomal adaptation dynamics. **a** To reflect the heterogeneous adaptation dynamics present in real bacterial cells, we can modify our original three-sector model to include multiple ribosomal subsectors, each with relative flux allocation *c*_*m*_ and characteristic relaxation timescale *λ*_*m*_. **b** Subsector flux allocation follows a power law with respect to relaxation timescale, specifically 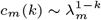, where the exponent *k* is the memory strength. Increases in memory correspond to increases in allocation to subsectors with faster relaxation timescales. **c**,**d** Example simulations of ribosomal subsector allocation fraction, *c*_*m*_(*k*), overall subsector allocation, 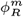, and resulting growth rate, *κ*, for a seven subsector model initialized in poor nutrients and the subjected to single nutrient upshift (**c**) or nutrient-rich pulsing (**d**). The environmental estimate used to determine *k*, is computed using the approximate implementation of the Bayesian inference scheme (Supplementary Note 2). In **d**, the bias for rich nutrients (*α*, see Methods for more details) is set to zero to clearly show how allocation is affected by nutrient pulsing. See Table 2 in Supplementary Information for a list of parameters.

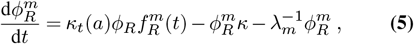

where 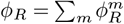and where 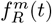 denotes the fractionof total synthesis capacity devoted to ribosomal subsector *m*. As before, these *m* equations are coupled to the dynamics of *a*(*t*) (Eq. (2)), yielding a system of *m*+1 coupled ODEs. The first term of Eq. (5) denotes the ribosomal production rate, the second captures dilution due to growth, and the third represents additional negative feedback control, with rate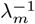. Importantly, in Eq. (5) when *λ*_*m*_ *≫* 0, then the third term vanishes and *κ*_*t*_(*a*) controls the timescale of adaptation for that sector, as in Eq. (1). In contrast, if *λ*_*m*_ is small such that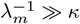, dilution due to growth no longer dominates, and inactivation (third term) dictates the relaxation dynamics. Simulating this model for a limited number of subsectors shows clearly how *λ*_*m*_ affects subsector dynamics, 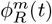, in dynamic nutrient environments. Subsectors with larger values of *λ*_*m*_ respond slower to nutrient perturbations than those with smaller values (Supplementary Fig. 2).

To describe how the cell dynamically regulates each subsector based on environmental context, we introduce *c*_*m*_(*t*), the fraction of the total translational flux allocated to subsector *m*, such that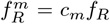. We relate this allocation to the subsector’s relaxation timescale *λ*_*m*_ and the cell’s internal estimate of environmental uncertainty (memory), *k*(*t*), using the relation: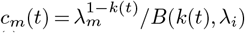, where 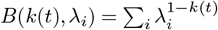 is a normalization constant to ensure ∑_*m*_ *c*_*m*_ (*t*) = 1 (see Supplementary Note 3 for detailed derivation). Thus, as the cell updates its internal representation of the environment through changes in *k*(*t*), it alters the relative weights *c*_*m*_(*t*), thereby re-allocating translation capacity across subsectors with different relaxation timescales (Fig. 4b).

Using our approximate Bayesian inference scheme and Eq. (5) along with our expression for *c*_*m*_(*t*), we can simulate the effect of different environments on subsector allocation. In response to a single upshift, allocation to subsectors with slower relaxation dynamics increases at the expense of subsectors with faster relaxation dynamics, indicative of the cell removing negative feedback control to maximize growth (Fig. 4c). In contrast, allocation to faster subsectors increases in fluctuating environments, allowing for rapid ribosomal, and thus growth rate adaptation (Fig. 4d).

In this formulation, the dynamics of the entire ribosomal sector can be approximated as (see Supplementary Note 3):

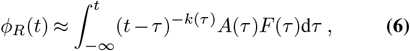

where *A*(*τ*) = *k*(*τ*)*b*^−*k*(*τ*)^Γ(*k*(*τ*)) and *b* is the maximum ribosomal inactivation rate. Remarkably, Eq. (6) shows that a model of multiple ribosomal subsectors with heterogeneous adaptation dynamics can give rise to overall ribosomal sector dynamics which contain a power law memory kernel of order *k*, thus recapitulating the key feature of our phenomenological nonlocal-in-time dynamical model (Eq. (3)). Critically, simulating this model for different nutrient pulse frequencies predicts an increase in effective adaptation rate with time period (Supplementary Fig. 3). Furthermore, this result allows us to use our mechanistic description to interpret the phenomenological power-law memory model. Specifically, the memory strength *k* can now be interpreted as a parameter describing the distribution of ribosomal relaxation timescales. Thus, the dynamic changes in memory *k* in fluctuating environments (Fig. 3a) can be interpreted to arise from an increase in allocation to ribosomal processes (subsectors) with faster relaxation timescales.

## Trade-off between growth rate maximization and adaptation speed

Introducing ribosomal subsectors has important consequences to the overall cellular growth rate, as inactivated ribosomes cannot contribute to growth. As a result, the total cellular growth rate is now given by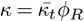, where

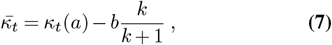

(see Supplementary Note 3). Here we see that 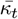 is the effective net translational efficiency, given by the weighted average of the translational efficiencies of each subsector. In the absence of inactivation (*b* = 0), we recover our original model for growth control with a single relaxation timescale. However, when negative feedback control is present, it is clear from Eq. (7) that increasing the proportion of ribosomes with fast relaxation dynamics (increasing *k*) *necessarily reduces* the overall growth rate of the cell (Fig. 5a). Thus Eq. (7) identifies an inherent tradeoff that cells must navigate between maximizing growth rate in the current environment (reducing *k*), and preserving the ability to adapt quickly to future changes in the environment (increasing *k*).

**Fig. 5.**
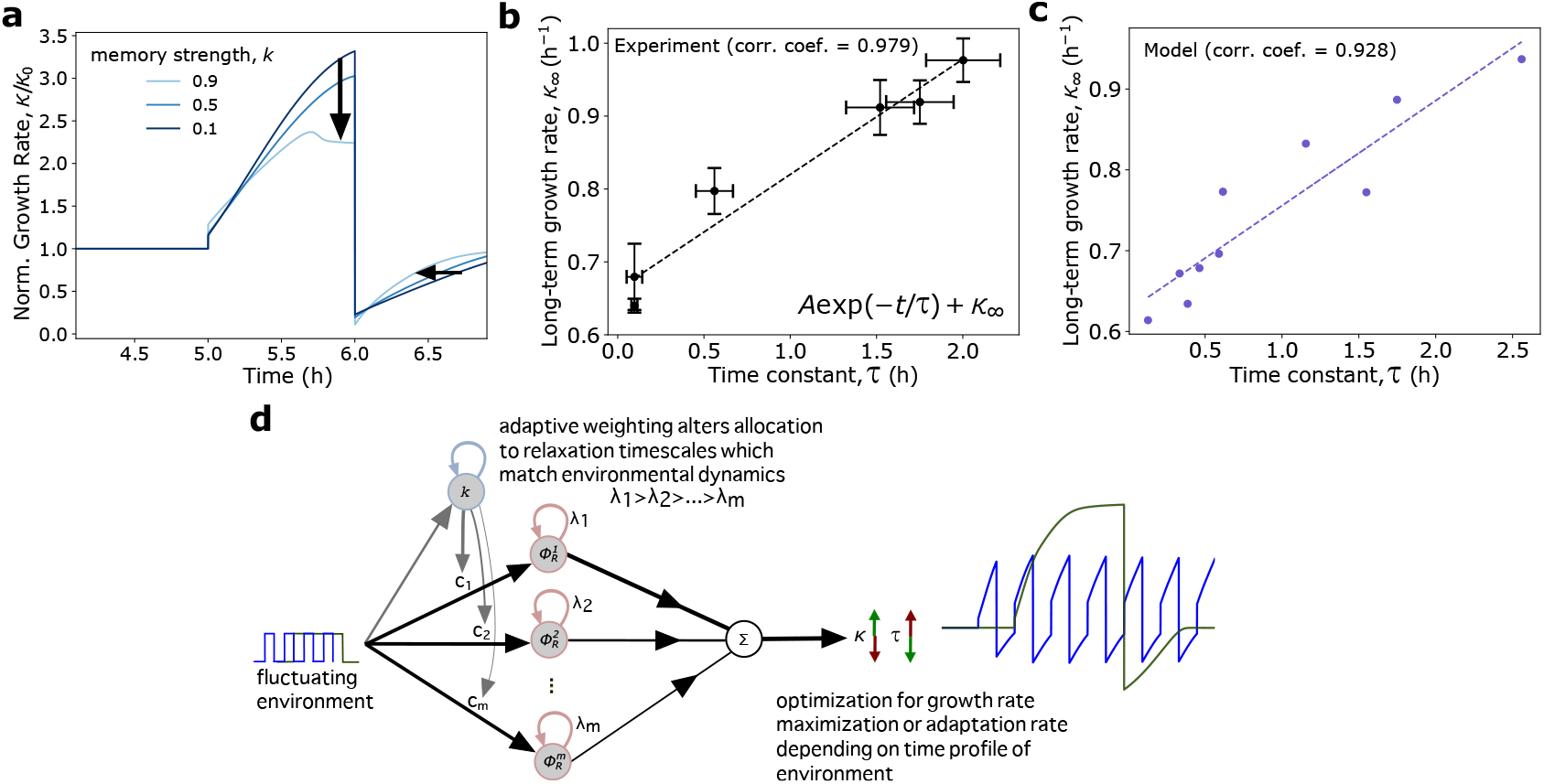
Mechanistic model predicts an inherent tradeoff between growth rate maximization and adaptation speed. **a** Example simulation of ribosomal subsector model with three subsectors with constant allocation for different memory strengths. Increasing memory reduces growth rate, but increases adaptation time. **b**,**c** Long-term growth rate, *κ*_*∞*_, exhibits a strong positive correlation with effective adaptation time constant, *τ*, in both experiment (**b**) and dynamic power-law memory model simulations (**c**). As with Fig. 1, *τ* and *κ*_*∞*_ were obtained by fitting growth rate data from the first nutrient-rich half-period after 6 h for different time periods to the exponential *A* exp(−*t/τ*) + *κ*_*∞*_, in order to maintain approximately the same amount of total nutrient exposure across all time periods. **d** Cells navigate the tradeoff between growth rate and adaptation by tuning subsector allocation in response to the time profile of nutrient fluctuations. See Table 1 in Supplementary Information for a list of parameters.

To test this prediction experimentally, we compared the relationship between the adaptation timescale, *τ*, and the long-term growth rate, *κ*_*∞*_, for different time periods of nutrient pulsing in both experiment and model simulations. Remarkably, we find a strong positive correlation between adaptation timescale and long-term growth rate for both cases (Fig. 5b,c), suggesting that growth is indeed limited by this constraint. This predicted trade-off has been observed experimentally in various organisms and settings (44–46), suggesting that this is a universal constraint governing growth in fluctuating environments, one which bacteria navigate through utilization of history-dependent resource allocation (Fig. 5d).

## Growth regulation as a recurrent neural network

Intriguingly, our mechanistic model can directly be mapped to continuous-time descriptions of artificial recurrent neural networks (RNNs) (47) (Fig. 6a). In this framing, the cellular reaction-network takes as input the external nutrient signal, *c*(*t*), and computes a corresponding growth rate, *κ*(*t*). Critically, this growth rate output is not only a function of *c*(*t*), but also of an internal or ‘hidden’ state vector comprised of subsector mass fractions **h**(*t*) = [*h*_1_, *h*_2_, …, *h*_*m*_]. Abstracting away the specific biological details, the evolution of each internal state can then generically be written as:

**Fig. 6.**
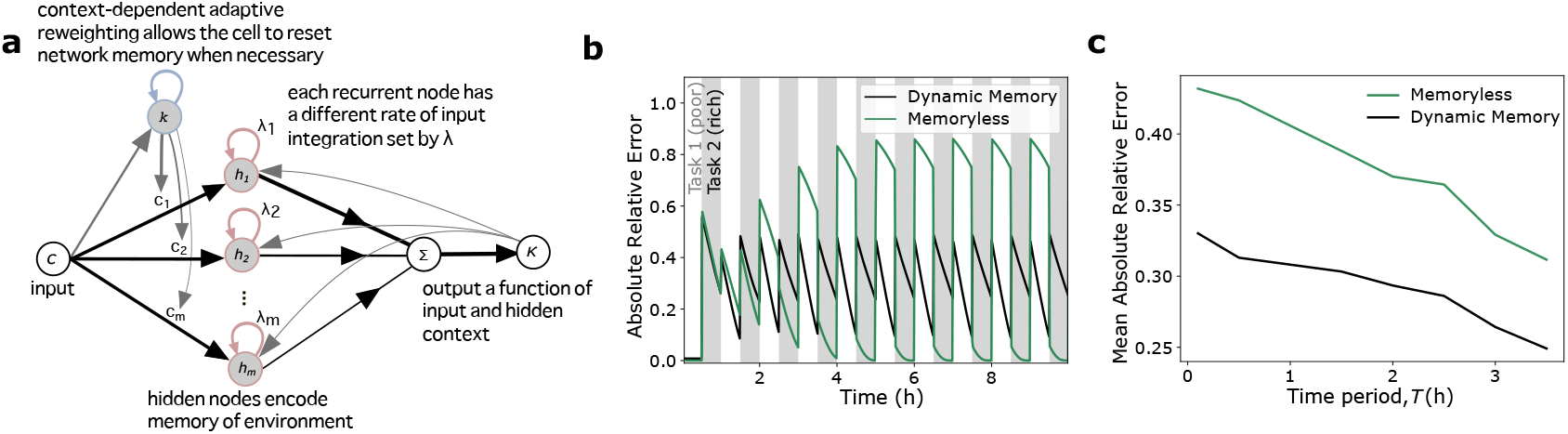
Recurrent reaction networks enable efficient continual learning in fluctuating environments. **a** The features enabling efficient task switching in the mechanistic (dynamic memory) model can be understood when viewing the reaction network as an RNN. **b** ‘Test error’, defined as the absolute relative difference between the current growth rate and the steady-state maximum, as a function of time for cells initialized in poor nutrients (trained for task 1) which then subsequently experience nutrient rich pulsing (switching between task 1 and task 2), for both the memoryless and dynamic memory models. **c** Quantification of the average test error for the setup described in (**b**) for different time periods of ‘task switching’, simulated for 24 h. The dynamic memory model outperforms the memoryless model for all time periods, indicative of its ability to efficiently ‘task switch’, when the environmental input changes. See Table 1 in Supplementary Information for a list of parameters.

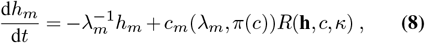

where *R*(·) denotes the recurrent input, which is a function not only of the hidden states **h**(*t*) but also the input *c*(*t*) and output feedback *κ*(*t*), and *c*_*m*_(·) denotes the subsector allocation.

Each hidden state (subsector) has a unique dynamic rate of input integration controlled by *λ*_*m*_ and output feedback from *κ*, together encoding a complex memory of the environment through subsector allocation, enableing rapid adaptation to previously seen environments. However, a network that maintains a strong memory of past inputs and cannot be reset when needed, such that its output remains stable regardless of changes to the input, is not very useful. This tradeoff between maintaining a memory of previous information and learning to adapt to new information has been identified in many contexts in both the machine learning and neuroscience literature (47–50). One way this tradeoff can be navigated by RNNs is by the inclusion of various gating mechanisms (47, 49), which allow for flexible updating or erasure of memory in a context-dependent manner. Remarkably, the control of subsector allocation, *c*_*m*_(·), by the cell’s internal representation of the environment, *π*(*c*), can be identified as an input gating mechanism which exactly serves this purpose. This simple gating mechanism exhibits functions similar to the input, forget, and output gates found in Long Short-Term Memory RNNs (49). Specifically, environment-dependent dynamic re-weighting of the input signal to the different subsectors allows the cell to regulate the effect of each subsector on setting the overall output, thus allowing the cell to maintain a flexible memory of previous environmental context which can easily be ‘forgotten’ if this context is no longer relevant.

The advantage of this network structure can be clearly seen if growth rate adaptation is viewed as a ‘continual learning’ problem, a classic machine learning paradigm in which models are trained on multiple tasks sequentially instead of simultaneously (50). In this framework, the ‘task’ can be viewed as maximizing growth rate in a given nutrient environment. Thus in fluctuating environments, a cell must constantly deal with task switching, where the objective periodically changes. In this setting, the dynamic memory model is able to outperform the memoryless model over a wide range of time periods by maintaining a low average error across both tasks (Fig. 6b,c). In contrast, the memoryless model is unable to efficiently task switch due to its lack of adaptive memory.

Together, this work demonstrates that despite their relative simplicity, single-celled organisms can exhibit complex learning behavior driven by adaptive memory encoded in intracellular reaction-networks. This learning behavior is critical in ensuring robust adaptation to stress in fluctuating environments.

## Methods

### Experimental data acquisition and image analysis. Experimental setup

With time-lapse microscopy, microfluidic mother machine devices (51, 52) allow accurate measurements for transient growth rates of single cells in preprogrammed microenvironments. PDMS devices were fabricated as described by Wang et al. (51). For each main trench on a PDMS device, two inlet holes were drilled to facilitate medium switching and oscillating. The PDMS device is then bound to a WillCo-dish (HBST-5040) glass bottom. Before use, machine devices are preprocessed in a plasma cleaner (Harrick Plasma PDC-32G) for 10 minutes at 0.8 Torr, baking at 80 °C for 10 minutes and primed with 0.5 mg/ml bovine serum albumin fraction V (BSA, Roche Diagnostics, CAS-No. 9048-46-8) for 10 minutes. Each main trench on the mother machine microfluidics contains 4000 individual channels. After cells were injected into the main trench, cells were loaded into channels by centrifuging the device. The sample size represents the number of single-cell division cycles measured from each mother machine experiment. Supplementary Fig. 4 illustrates the configuration of the mother machine and syringe pump.

Fresh media was infused through the microfluidic device by a syringe pump after the cell loading. Syringe pumps (New-Era pump systems, SKU:4000) were programmed to pump sodium acetate (slow growth) medium for an initial 5 hours, and then pulse the slow growth medium and fast growth medium (glucose + 11 a.a.) with the set period of 1, 2, 3, 4, and 6 hour cycle time. The syringe pulsing program for our experiments can be found in Supplementary Table 3.

### Cell preparation

Cells from *E. coli* strain K-12 MG1655 background (FS003) (53) were used in this study. Prior to each time-lapse imaging experiment, cells were picked from glycerol stock in -80 °C and spread on an agar plate. After overnight culture at 30 °C, cells were inoculated into 1 ml LB medium (MP Biomedicals, LLC, SKU: 113002122-CF). The cultures were then shaken for 12-18 hours at 30 °C in a water bath shaker. Cell cultures were diluted into 2 ml of MOPS medium with sodium acetate, and shaken in the water bath at 37 °C for 24 hours. The cultures were then concentrated 10-to 100-fold and injected into the microfluidic mother machine device. Fresh media was infused with a syringe pump through the duration of the experiment. See Supplementary Tables 4, 5, and 6 for full media description and details.

### Image acquisition

We performed phase-contrast imaging on a Nikon ECLIPSE Ti2 inverted microscope with 100x oil immersion objective (PH3, numerical aperture = 1.45), and ORCA-Fusion BT Digital CMOS camera (C15440-20UP). The syringe pump programs controlled medium pulsing with a pump rate of 0.8 ml/h for all experiments. Exposure time for phase-contrast imaging was set to 40 ms, and images were captured every two minutes.

### Image processing and data analysis

Image processing followed a previously developed mm3 software pipeline (52, 54). The mm3 image analysis pipeline includes channel compilation and designation, background subtraction, cell segmentation, cell tracking through lineages, and data output and analysis. Single-cell physical properties, such as length, width, and growth rates were calculated throughout the entire timecourse of the experiments. To calculate the instantaneous growth rate, *κ*, we considered single-cell growth an exponential process and fit an exponential to a sliding window of three data points. The resulting growth rates were then binned in 0.05 h intervals and averaged across the population.

### Updating memory strength through a Bayesian inference scheme

To connect memory strength *k* to the nutrient environment *c*, utilize a Bayesian inference scheme in which the cell dynamically updates its estimate of the environmental uncertainty based on previous environmental signals. For pulsatile nutrient exposure, we can simply describe the environment as fluctuating between two nutrient states *X*_*i*_ *∈* {0, 1}, rich (1) and poor (0). In a discrete setting, the signal history describing such a fluctuating environment can be written as a sum *E*_*n*_ = *X*_1_ + *X*_2_ + … + *X*_*n*_ of *n* independent *X*_*i*_ ∼ Bernoulli(*π*) random variables, where *π* parameterizes the environment. The likelihood of receiving *m* nutrient-rich signals over *n* observations is then:

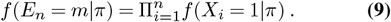

Inserting the functional form, *f* (*x*| *π*) = *π*^*x*^(1 − *π*)^1−*x*^, this is then:

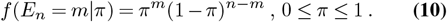

To adjust memory strength to an appropriate level, the cell must estimate the statistics of the environment based on the most recent observations. This can be obtained by maximizing the posterior of the parameters conditioned on the observations, *f* (*π*|*E*_*n*_). Using Bayes’ rule, that is:

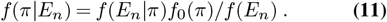

Naturally, past information can be incorporated using the conjugate prior for a sequence of Bernoulli observations, which is a beta distribution:

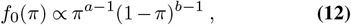

thus our posterior distribution becomes:

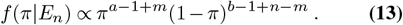

For a uniform prior *f*_0_ = 1, in which we do not impose any historical belief about the environment, then *a* = *b* = 1, and the maximum a posteriori probability (MAP) equals the maximum likelihood estimate (MLE), which is simply given by *π* = *m/n*. However, given a new observation *X*_*n*+1_, the posterior distribution and corresponding MAP can be updated by revising the current belief with the new likelihood, specifically:

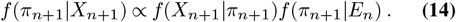

For a fixed recall history *N* starting at time *t*, this finally yields the iteratively updating distribution:

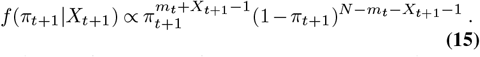

As before, the corresponding MAP is simply given by *π*_*t*_ = *m*_*t*_*/N*, thus it is sufficient for the cell to track the number of observed nutrient-rich signals *m*_*t*_ over the last *N* measurements.

A cell’s gene regulatory networks may encode a bias to expect certain environments. Previous work has suggested that cells maintain an excess of biosynthetic pathways to quickly take advantage of nutrient upshifts (2). This type of persistent bias can be incorporated into our Bayesian inference scheme via a mixture prior that combines a fixed bias component with the iteratively updating prior:

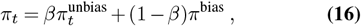

where *β* controls the strength of the influence of the bias. If the cell is biased to expect rich nutrients, such that *π*^bias^ = 1, this simplifies to:

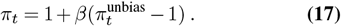

This approach is equivalent to adding *α* rich-nutrient signal pseudocounts at each time step. This can be seen by letting *β* = *N/*(*N* + *α*), such that *π*_*t*_ becomes:

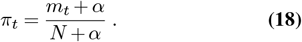

In a continuous environment, such as the one we model in this work, this is equivalent to tracking the moving average of the nutrient signal, 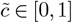, over the interval [*t* − *T*_*N*_, *t*]:

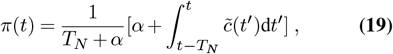

where here now the bias parametrized by *α* is in units of time. Critically, this estimate of environmental uncertainty introduces a new timescale into the dynamics set by *T*_*N*_, which determines the rate at which new observations reshape the cell’s estimate of the environment. Specifically, a larger value of *T*_*N*_ means the cell takes more history into account when making an estimate of the environment, allowing for more accurate estimates but at the cost of responding slower to changes in the underlying distribution. In Supplementary Note 2 we show how this type of computation could approximately be carried out by the cell.

In maximally uncertain environments (when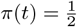), the timescale of growth rate adaptation, and thus proteome reallocation, should be minimized to allow for rapid adaptation to unpredictable environmental changes. Thus we relate the memory strength at time *t, k*(*t*), to the MAP at time *t, π*(*t*), through a simple linear relation, specifically:

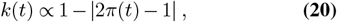

such that *k* increases in fluctuating environments. Importantly, if a bias is present, *k* = 0 regardless of *α* in constant nutrient-rich conditions, enabling growth rate maximization. However, in constant nutrient-poor conditions, *α >* 0 ensures that memory is also nonzero, as *π* = *α/*(*T*_*N*_ + *α*), allowing for faster adaptation in response to nutrient upshift.

### Numerical Simulations

Numerically integrating FDEs are more complex than their ODE counterparts due to the time nonlocality present in the dynamics. To simulate the variable, multi-order system of FDEs which define our model (Eq. (2) and Eq. (3)), we extended a previously-developed predictor-corrector method for constant-order FDEs based on Adams formulae (55, 56) to the variable-order case. Our fractional model can be expressed as an initial value problem:

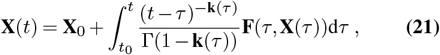

where **X**(*t*) = [*ϕ*(*t*), *a*(*t*)], **k**(*t*) = [*k*(*t*), 0], and **F**(*t*) = [*F* (*t*), *J*(*t*)] where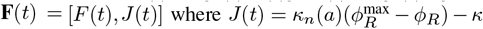. To solve Eq. (21) we employed a product integration technique (56). Briefly, for grid nodes *t*_*j*_ (where *j* = 0, …, *m*) with constant step size *h*, first the explicit product rectangle rule gives a prediction for Eq. (21) at *t*_*j*_:

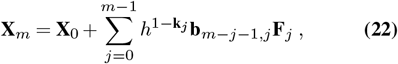

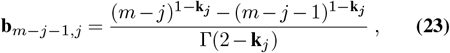

where **F**_*j*_ = **F**(*t*_*j*_, **X**_*j*_) and **X**_*j*_ denotes the numerical approximation to **X**(*t*_*j*_). Second, the implicit product trapezoidal rule provides a corrector estimate of Eq. (21):

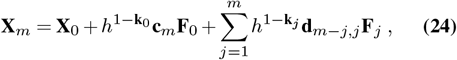

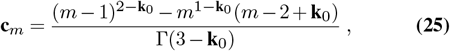

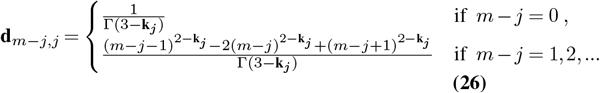

The last term of Eq. (24), **F**_*m*_ = **F**(*t*_*m*_, **X**_*m*_), is calculated using the value for **X**_*m*_ obtained using Eq. (22). Computing this algorithm is costly, as evaluation of the solution on the entire grid *t*_0_, *t*_1_, …, *t*_*N*_ is proportional to 𝒪 (*N* ^2^) (56).

## Acknowledgments

S.B. acknowledges support from the National Institutes of Health (NIH R35 GM143042), and the Shurl and Kay Curci Foundation. J.C.K. acknowledges support from the National Institutes of Health (NIH T32 GM133353). F.S. acknowledges support from the National Science Foundation (NSF/MCB-BSF:2309595).

## Author Contributions

J.C.K. and S.B. designed and developed the theory. H.W. and F.S. designed the experiments. J.C.K. performed the model simulations and analyzed the data. H.W. performed the experiments and image analysis. J.C.K. and S.B. drafted the paper. J.C.K., H.W., F.S and S.B. edited the paper.

