## Supplementary Information for "Power-law memory governs bacterial adaptation and learning in fluctuating environments"

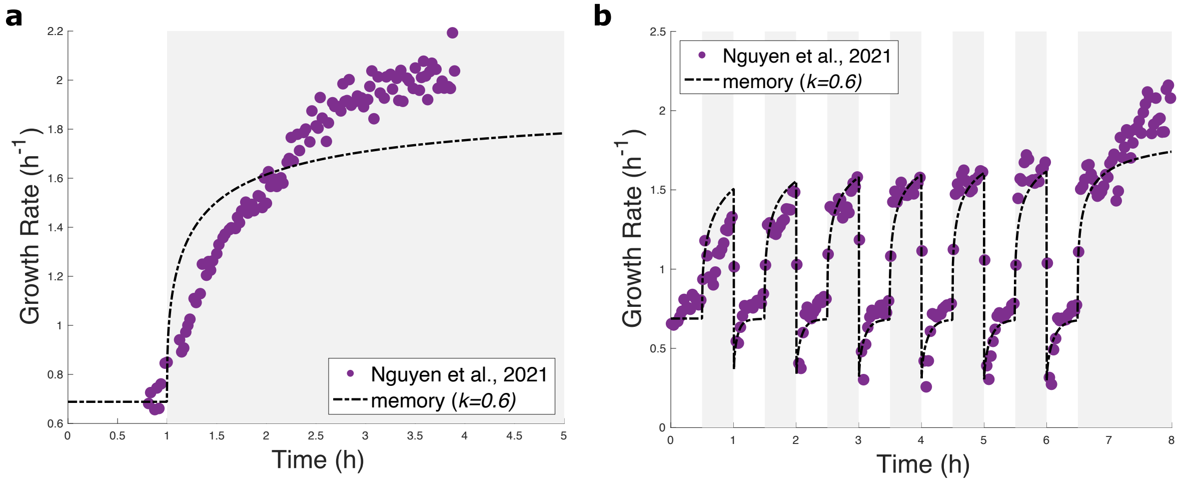


Figure 1: **Constant-order fractional model unable to capture observed single-shift and pulsatile growth rate dynamics simultaneously**. Resulting growth control dynamics for single nutrient upshift (**a**) and nutrient pulsing (**b**) for a constant memory strength value of $k = 0.6$. Memory strength value was obtained through fitting to the pulsatile nutrient data (**b**), and then simulating the resulting single shift dynamics (**a**). Although the constant-order model does a good job of capturing pulsatile data, it is unable to capture the single-shift response using the same parameter values – motivating a variable-order fractional formulation. Experimental data are from ref. [1] and are of *E. coli* K-12 NCM3722 $\Delta motA$ cells switching between low (0.1%) and high (2%) concentrations of LB.

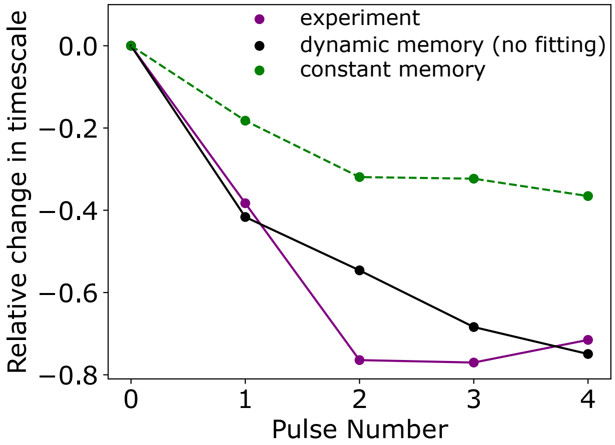


Figure 2: **Constant-order model unable to capture episodic learning behavior**. Relative change in adaptation timescale $\tau$ for cells initially grown in nutrient-poor media and exposed to subsequent nutrient-rich pulses. Adaptation timescale decreases before stabilizing in both experiment and in the dynamic power-law memory model. The constant-order memory model is unable to capture the same magnitude decrease. Experimental data corresponds to that shown in Fig. 3a,b in the main text, and are of *E. coli* K-12 MG1655 cells subjected to nutrient shifts between MOPS sodium acetate and MOPS glucose with amino acids using a custom microfluidic device (see Methods).

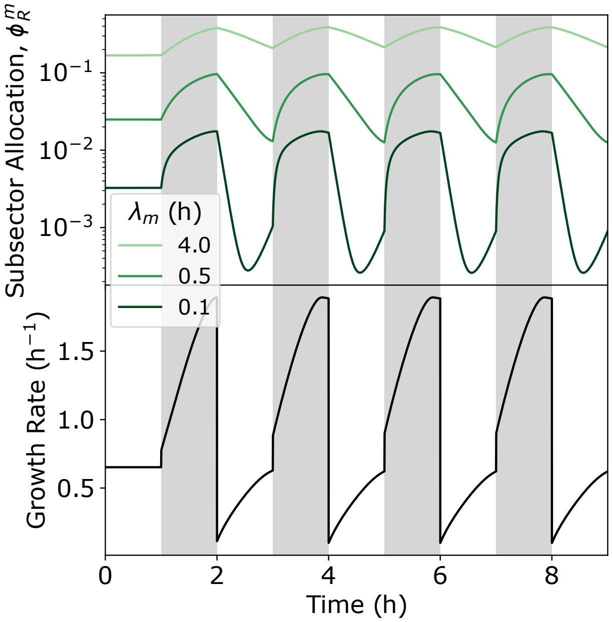


Figure 3: **Subsectors with larger values of $\lambda_m$ respond slower to nutrient perturbations than those with smaller values**. Ribosomal subsector allocation, $\phi_R^m$, and corresponding growth rate dynamics for a three subsector model with a constant value of $k = 0.6$.

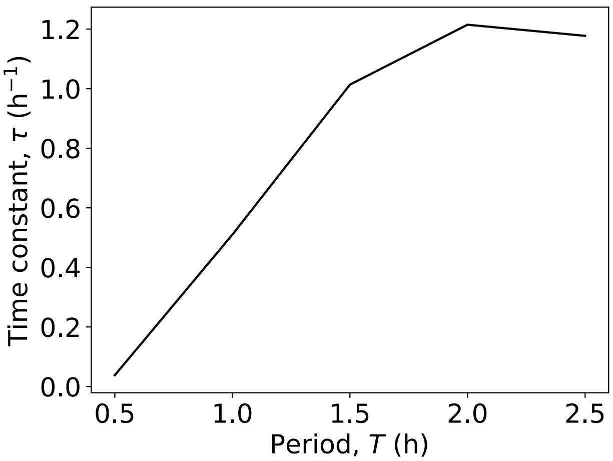


Figure 4: **Ribosomal subsector model captures the increase in adaptation rate with time period of nutrient pulsing seen experimentally and in the variable-order fractional model**. As before, for each time period of nutrient pulsing, $T$, the effective adaptation time constant, $\tau$, was obtained by fitting the first upshift after 6 h to the exponential $A\exp(-t/\tau) + \kappa_\infty$.

### 1 Derivation of minimal model

To initially predict cellular growth rate, we utilized a simplified version of our previously published framework [2]. Our minimal model captures dynamic proteome allocation to three coarse-grained sectors consisting of housekeeping proteins (Q), ribosomal proteins (R), and metabolic proteins (P). Bacterial cells grow exponentially in size during the cell cycle. Assuming constant protein density, the growth rate $\kappa$ of a single cell can be defined in terms of cell volume, $V$, or equivalently in terms of total protein mass, $M$, yielding

$$\kappa = \frac{1}{M}\frac{\mathrm{d}M}{\mathrm{d}t} = \frac{1}{V}\frac{\mathrm{d}V}{\mathrm{d}t} \ . \tag{1}$$

The rate of change of protein mass is proportional to the mass of actively translating ribosomes. Thus, the rate of change of protein mass is given by

$$\frac{\mathrm{d}M}{\mathrm{d}t} = \kappa_t M_R \ , \tag{2}$$

where $\kappa_t$ is the translational efficiency of the cell and $M_R$ is the total mass of ribosomes. Using Eqs. (1) and (2), the growth rate can then be defined as $\kappa = \kappa_t \phi_R$, where $\phi_R = M_R/M$ is the ribosome mass fraction, thus recovering Eq. (1) in the main text. To obtain the dynamics of $\phi_R$, we note that $\frac{\mathrm{d}M_R}{\mathrm{d}t} = \kappa_t f_R M_R$, where $f_R$ is the fraction of total cellular protein synthesis flux devoted to ribosomes. It follows that the time dynamics of $\phi_R$ are then

$$\frac{\mathrm{d}\phi_R}{\mathrm{d}t} = \kappa_t(a)\phi_R(f_R(a) - \phi_R) \ , \tag{3}$$

thus recovering Eq. (2) in the main text. As noted previously, both $\kappa_t$ and $f_R$ depend on the intracellular amino acid concentration, which in turn depends on the nutrient availability. To connect cellular growth rate to amino acid mass fraction ($a$) and nutrient availability ($c$), we use the following condition for flux balance [3]: $\frac{\mathrm{d}a}{\mathrm{d}t} = \kappa_n(a)\phi_P - \kappa$, where $\kappa_n$ is the nutritional efficiency of the cell and $\phi_P$ is the mass fraction of P-sector protein that are responsible for transporting nutrients into the cell. Using the constraint $\phi_R + \phi_P = 1 - \phi_Q = \phi_R^{\max}$, along with our definition of growth rate from Eq. (1), the amino acid mass fraction can be rewritten as

$$\frac{\mathrm{d}a}{\mathrm{d}t} = \kappa_n(a)(\phi_R^{\max} - \phi_R) - \kappa \ , \tag{4}$$

finally recovering Eq. (3) in the main text. To make explicit the dependency of the efficiencies, $\kappa_n$ and $\kappa_t$, on $a$, we define two regulatory functions, $f(a)$ and $g(a)$, as given by [4]. Specifically, we assume $\kappa_n = \kappa_n^0(c)f(a)$ and $\kappa_t = \kappa_t^0 g(a)$, where $\kappa_t^0$ is a constant, and $\kappa_n^0$ is a function of the extracellular nutrient concentration $c$. The regulatory functions are then

$$f(a) = \frac{1}{1 + (a/a_n)^2} \ , \tag{5}$$

$$g(a) = \frac{(a/a_t)^2}{1 + (a/a_t)^2} \ , \tag{6}$$

where translation becomes significantly attenuated for amino acid concentrations below $a_t$, and the amino acid supply flux becomes significantly attenuated by feedback inhibition for $a$ above $a_n$. Finally, $f_R$ is chosen to maximize amino acid flux, and equivalently growth rate, at steady state [2]. This yields the functional dependence of $f_R$ on amino acid concentration

$$f_R(a) = \frac{-f'(a)g(a)\phi_R^{\max}}{-f'(a)g(a) + f(a)g'(a)} \ , \tag{7}$$

where $f'$ and $g'$ denote the derivative of $f$ and $g$ w.r.t. $a$, respectively.

### 2 Approximate implementation of a Bayesian inference scheme through cellular reaction networks

In the Methods section in the main text we present a Bayesian inference scheme to connect the time-profile of nutrient fluctuations to memory strength. Here we show how such a scheme could be implemented by the cell.

For a fixed recall of length $N$ and nonexchangeable prior $\alpha$, the corresponding iterative updating scheme given by the MAP is $\pi_t = \beta\pi_t^{\text{unbias}} + (1-\beta) = (m_t+\alpha)/(N+\alpha)$, where $\pi_t^{\text{unbias}}$ denotes the unbiased cell estimate. $\pi_t^{\text{unbias}}$ is simply given by the fraction of nutrient-rich signals, $m_t$, observed over the past $N$ observations starting from time $t$:

$$\pi_t^{\text{unbias}} = \frac{m_t}{N} = \frac{\sum_{i=t-N-1}^{t} X_i}{N} \ , \tag{8}$$

where $X_i \in 0,1$ denotes the external nutrient environment being either rich (1) or poor (0) at time $i$. To compute this moving average requires keeping track of the last $N$ environmental signals. However, the cell can approximate this value by instead keeping an exponential moving average of past nutrient signals – an approach which only requires storage of the most recent estimate. To see this, we can equivalently write $\pi_t^{\text{unbias}}$ starting at $t=1$ as:

$$\pi_N^{\text{unbias}} = \frac{X_N + \sum_{i=1}^{N} X_i}{N} \ , \tag{9}$$

$$= \frac{N-1}{N}\frac{\sum_{i=1}^{N-1} X_i}{N-1} + \frac{X_N}{N} \ . \tag{10}$$

The summation $\sum_{i=1}^{N-1} X_i/(N-1)$ represents the fraction of nutrient-rich signals over the past $N-1$ observations, starting at $t=N-1$. If $N >> 1$, this is approximately equal to the sum $\pi_{N-1}^{\text{unbias}} = \sum_{i=0}^{N-1} X_i/N$, thus we can approximate $\pi_t^{\text{unbias}}$ as:

$$\pi_N^{\text{unbias}} = (1-\frac{1}{N})\pi_{N-1}^{\text{unbias}} + \frac{X_N}{N} \ . \tag{11}$$

This result now gives us an iterative updating scheme which only requires memory of the previous estimate:

$$\pi_t^{\text{unbias}} = (1-\rho)\pi_{t-1}^{\text{unbias}} + \rho X_t \ , \tag{12}$$

where here $\rho = 1/N$ is often referred to as the *learning rate*, and sets the timescale over which new data reshape the cell's estimate of the environment.

The finite difference method can be used to convert Eq. (12) to continuous time, yielding:

$$\frac{\mathrm{d}\pi^{\text{unbias}}(t)}{\mathrm{d}t} \approx \frac{\pi_t^{\text{unbias}} - \pi_{t-1}^{\text{unbias}}}{\Delta t} = \tilde{\rho}(\tilde{c}(t) - \pi^{\text{unbias}}(t)) \ , \tag{13}$$

where $\tilde{\rho} = \rho/\Delta t$ and $\tilde{c}(t)$ is the continuous analog of $X_t$. Integrating Eq. (13), we arrive at an expression for $\pi^{\text{unbias}}(t)$:

$$\pi^{\text{unbias}}(t) = \tilde{\rho}\int_{-\infty}^{t} e^{-\tilde{\rho}(t-\tau)}\tilde{c}(\tau)\mathrm{d}\tau \ . \tag{14}$$

Thus, the cell can compute an approximate estimate of the environment through leaky integration of the nutrient signal. Here, $\tilde{\rho}$ sets the leak rate, thus controlling how quickly the cell updates its estimate. This type of computation can easily be implemented by the cell via a multi-step reaction pathway which is a function of the environmental nutrient signal $\tilde{c}(t)$ [5].

### 3 Derivation of power law memory kernel from ribosomal subsector model with diverse relaxation timescales

Here we show how a fractional model of growth control can emerge from a model of ribosomal subsectors with varying relaxation timescales. Let the ribosomal mass fraction, $\phi_R$, be composed of $m$ subsectors, each with a unique relaxation timescale $\lambda_m$. Modifying Eq. (3) to incorporate these changes yields the dynamics:

$$\phi_R^m + \lambda_m\frac{d\phi_R^m}{dt} = \lambda_m\kappa_t(a)\phi_R f_R^m(a) - \lambda_m\phi_R^m\kappa \ , \tag{15}$$

where

$$\phi_R = \sum^m \phi_R^m \ . \tag{16}$$

Dynamic subsector reallocation is driven by resource allocation strategies encoded by gene-regulatory networks which depend on the external nutrient environment. To account for these effects, let $c_m(t)$ and $\gamma_m(t)$ denote the proportion of the total ribsomal sector allocation and translational flux comprising subsector $m$, respectively. This substitution yields:

$$\phi_R^m + \lambda_m \frac{d\phi_R^m}{dt} = \lambda_m \kappa_t(a) \phi_R c_m(t) f_R(a) - \lambda_m \gamma_m(t) \phi_R \kappa \ . \tag{17}$$

Assuming that subsector reallocation can occur by a variety of mechanisms (not solely requiring the production of new ribosomes), such that allocation can equilibrate much faster than the dynamics of the full sector $\phi_R$, we use the quasi-steady-state approximation to obtain the relationship between $c_m$ and $\gamma_m$: $\gamma_m = \frac{\kappa_t(a) f_R(a)}{\kappa + \lambda_m^{-1}} c_m$. For most sectors $\lambda_m^{-1} << \kappa$ making $\kappa_t(a) f_R(a) \approx \kappa$, thus we take $c_m(t) \approx \gamma_m(t)$, and arrive at an expression for $\phi_R^m$ in terms of the total net production-dilution flux, $F(t) = \phi_R(\kappa_t(a) f_R(a) - \kappa)$, specifically:

$$\phi_R^m + \lambda_m \frac{d\phi_R^m}{dt} = \lambda_m c_m(t) F(t) \ , \tag{18}$$

As before, these $m$ equations are coupled to the dynamics of $a$ given by Eq. (4), yielding a system of $m+1$ coupled ODEs.

To obtain an expression for the time evolution of the entire ribosomal sector, $\phi_R(t)$, we start by multiplying Eq. (18) through by $\frac{1}{\lambda_m} e^{t/\lambda_m}$:

$$\frac{1}{\lambda_m} e^{t/\lambda_m} \phi_R^m + e^{t/\lambda_m} \frac{d\phi_R^m}{dt} = c_m(t) e^{t/\lambda_m} F(t) \ , \tag{19}$$

Making use of the chain rule, this is then:

$$\frac{d}{dt}[e^{t/\lambda_m} \phi_R^m] = c_m(t) e^{t/\lambda_m} F(t) \ , \tag{20}$$

Integrating over all past times until the time of interest $t$ and remembering that $\phi_R = \sum^m \phi_R^m$, an expression for the overall sector dynamics, $\phi_R$, can finally be obtained:

$$\phi_R = \int_{-\infty}^{t} (\sum^m c_m(\tau) e^{-(t-\tau)/\lambda_m}) F(\tau) d\tau \ . \tag{21}$$

We can relate allocation to each subsector, $c_m(t)$, to its relaxation timescale , $\lambda_m$, through a power law relation such that $c_m(t) = \lambda_m^{1-k(t)} / B(k(t), \lambda_i)$, where $B(k(t), \lambda_i) = \sum^i \lambda_i^{1-k(t)}$ is a normalization constant to ensure $\sum^m c_m(t) = 1$. We choose this specific relationship because the allocation distribution peaks at zero as a function of inactivation rate (as the most abundant subsector should be the one under no active negative feedback control), and it has a long tail to account for multiple mechanisms of feedback. This choice yields:

$$\phi_R = \int_{-\infty}^{t} (\sum^m \frac{\lambda_m^{1-k(\tau)}}{B(k(\tau), \lambda_i)} e^{-(t-\tau)/\lambda_m}) F(\tau) d\tau \ . \tag{22}$$

We note that now the time dependence of $c_m$ is through the exponent $k(t)$. If $m$ is large and $\lambda_m^{-1}$ sufficiently spaced, we can approximate this sum as an integral to give:

$$\phi_R = \int_{-\infty}^{t} (\int_a^b \frac{\lambda^{1-k(\tau)}}{B(k(\tau), a, b)} e^{-(t-\tau)/\lambda} d\lambda^{-1}) F(\tau) d\tau \ , \tag{23}$$

where now the normalization constant is a function of the bounds of integration, i.e. $B(k(t), a, b) = \int_a^b \lambda^{1-k(t)} d\lambda^{-1}$. For $a = 0$ and a maximum inactivation rate $b$, this becomes

$$\phi_R = \int_{-\infty}^{t} k(\tau) b^{-k(\tau)} (\Gamma(k(\tau)) - \Gamma(k(\tau), b(t-\tau)))(t-\tau)^{-k(\tau)} F(\tau) d\tau \ , \tag{24}$$

where $\Gamma(k(\tau))$ and $\Gamma(k(\tau), b(t-\tau))$ denote the gamma function and the upper incomplete gamma function, respectively. Thus, Eq. (24) shows that our model of ribosomal subsectors with diverse relaxation timescales gives rise to nonlocal-in-time dynamics for the overall $\phi_R$ sector, specifically yielding a memory kernel with power law decay of order $k$. When $t-\tau$ is large, $\Gamma(k(\tau)) \approx \Gamma(k(\tau)) - \Gamma(k(\tau), b(t-\tau))$, thus yielding the final expression we present in the main text:

$$\phi_R(t) \approx \int_{-\infty}^{t} k(\tau) b^{-k(\tau)} \Gamma(k(\tau)) (t-\tau)^{-k(\tau)} F(\tau) \mathrm{d}\tau \ . \tag{25}$$

Introducing ribosomal subsectors has important consequences to the overall cellular growth rate, as inactivated ribosomes cannot contribute to growth. As a result, the total cellular growth rate is now given by the summation of the net growth contribution from each subsector, yielding:

$$\kappa = \sum^{m} (\kappa_t(a) - \lambda_m^{-1}) \phi_R^m \ , \tag{26}$$

where here we omit the explicit time dependence for notational clarity. Approximating the sum as an integral and using our previous definition of $c_m$ we finally obtain:

$$\kappa = \bar{\kappa_t} \phi_R \ , \tag{27}$$

where

$$\bar{\kappa_t} = \kappa_t(a) - b \frac{k}{k+1} \ . \tag{28}$$

We present Eqs. (27) and (28) in the main text.

Table 1: Dynamic memory FDE model parameters.

| Parameter | Description | Value | Figure number |
|---|---|---|---|
| $\phi_R^{\max}$ | maximum flux allocation to ribosome production [3] | 0.55 | all |
| $a_t$ | translation attenuation threshold [4] | $10^{-4}$ | all |
| $a_n$ | feedback inhibition threshold [4] | $10^{-3}$ | all |
| $\kappa_t^0$ ($\text{h}^{-1}$) | translational efficiency rate constant, strain specific, fitted | 4.05 | 2 |
| | | 1.85 | 3,5 |
| $\kappa_{n,low}^0$ ($\text{h}^{-1}$) | nutritional efficiency rate constant in nutrient-poor media, calculated | 2.4 | 2 |
| | | 0.96 | 3,5 |
| $\kappa_{n,high}^0$ ($\text{h}^{-1}$) | nutritional efficiency rate constant in nutrient-rich media, calculated | 1500 | 2 |
| | | 21.59 | 3,5 |
| $T_N$ (h) | time window used to calculate MAP in Bayesian inference scheme, fitted | 1.9 | 2 |
| | | 1.30 | 3,5 |
| $\alpha$ (h) | bias parameter for rich nutrients used to calculate MAP in Bayesian inference scheme, fitted | 0 | 2 |
| | | 0.50 | 3,5 |
| $S$ | proportionality constant connecting $k$ to $\pi$ in Eq. (6) in the main text, fitted | 0.70 | 2 |
| | | 0.40 | 3,5 |

Table 2: Ribosomal subsector model parameters.

| Parameter | Description | Value |
|---|---|---|
| $\phi_R^{\max}$ | maximum flux allocation to ribosome production [3] | 0.55 |
| $a_t$ | translation attenuation threshold [4] | $10^{-4}$ |
| $a_n$ | feedback inhibition threshold [4] | $10^{-3}$ |
| $\kappa_t^0$ ($\text{h}^{-1}$) | translational efficiency rate constant | 4.5 |
| $\kappa_{n,low}^0$ ($\text{h}^{-1}$) | nutritional efficiency rate constant in nutrient-poor media | 2.5 |
| $\kappa_{n,high}^0$ ($\text{h}^{-1}$) | nutritional efficiency rate constant in nutrient-rich media | 70 |
| $\tilde{\rho}$ | learning rate (Eq. (14)), analogous to $T_N$ in the dynamic memory FDE model | 1 |
| $\beta$ | bias parameter for rich nutrients (Eq. (20) in the main text) | 0.6 |
| $S$ | proportionality constant connecting $k$ to $\pi$ (Eq. (6) in the main text) | 1 |

Table 3: Experimental conditions and sample size.

| | Experiment type | Pulse period (h) | # of pulsing cycles | Post Pulsing (h) | Sample size |
|---|---|---|---|---|---|
| 1 | Nutrient pulse | 1 | 12 | 12 | 29401 |
| 2 | | 2 | 6 | 12 | 30476 |
| 3 | | 3 | 6 | 8 | 29724 |
| 4 | | 4 | 6 | 8 | 51545 |
| 5 | | 6 | 6 | 8 | 22160 |
| 6 | Nutrient upshift | - | - | 8 | 3299 |

Table 4: Media Composition Table

| **Media Name (as used in the text)** | **Buffer** | **Carbon Source (v/w) Concentration** |
|---|---|---|
| Glucose + 11 a.a. | MOPS modified buffer | Glucose 0.2% |
| Sodium acetate | MOPS modified buffer | Sodium acetate 60 mM |

Table 5: MOPS modified buffer components.

| # | **Components** | **Concentration** |
|---|---|---|
| 1 | MOPS (MW 209.3) | 40 mM |
| 2 | Tricine (MW 179.17) | 4 mM |
| 3 | Iron(III) sulfate (MW 278.01) | 0.1 mM |
| 4 | Ammonium chloride | 9.5 mM |
| 5 | Sodium sulfate | 0.276 mM |
| 6 | Calcium chloride | 0.5 $\mu$M |
| 7 | Magnesium chloride | 0.525 mM |
| 8 | Sodium chloride | 50 mM |
| 9 | Ammonium molybdate | 3 nM |
| 10 | Boric acid | 0.4 $\mu$M |
| 11 | Cobalt chloride | 30 nM |
| 12 | Cupric sulfate | 10 nM |
| 13 | Manganese chloride | 80 nM |
| 14 | Zinc sulfate | 1 nM |
| 15 | Potassium phosphate monobasic | 1.32 mM |

Table 6: Supplements for Glucose + 11a.a.

| | **Components** | **Concentration ($\mu$g/ml)** |
|---|---|---|
| 1 | L-methionine (M) | 500 |
| 2 | L-histidine (H) | 500 |
| 3 | L-arginine (R) | 500 |
| 4 | L-proline (P) | 500 |
| 5 | L-threonine (T) | 500 |
| 6 | L-tryptophan (W) | 500 |
| 7 | L-leucine (L) | 500 |
| 8 | L-tyrosine (Y) | 500 |
| 9 | L-alanine (A) | 500 |
| 10 | L-asparagine (N) | 500 |
| 11 | L-aspartic acid (D) | 25 |
